# Interactions between metabolism and growth can determine the co-existence of *Staphylococcus aureus* and *Pseudomonas aeruginosa*

**DOI:** 10.1101/2022.09.14.507888

**Authors:** Camryn Pajon, Marla C. Fortoul, Gabriela Diaz-Tang, Estefania Marin Meneses, Taniya Mariah, Brandon Toscan, Maili Marcelin, Allison J. Lopatkin, Omar Tonsi Eldakar, Robert P. Smith

**Affiliations:** Department of Biological Sciences, Halmos College of Arts and Sciences, Nova Southeastern University, Fort Lauderdale FL, 33314; Department of Biology, Barnard College, Columbia University, New York NY, USA; Data Science Institute, Columbia University, New York NY, USA; Department of Ecology, Evolution, and Environmental Biology, Columbia University, New York NY, USA; Cell Therapy Institute. Kiran Patel College of Allopathic Medicine, Nova Southeastern University, Fort Lauderdale FL, 33314

**Keywords:** metabolism, polymicrobial community, ATP, growth rate, co-culture

## Abstract

Most bacteria exist and interact within polymicrobial communities. These interactions produce unique compounds, increased virulence and augmented antibiotic resistance. One community associated with negative healthcare outcomes consists of *Pseudomonas aeruginosa* and *Staphylococcus aureus*. When co-cultured, virulence factors secreted by *P. aeruginosa* reduce metabolism and growth in *S. aureus*. When grown *in vitro* this allows *P. aeruginosa* to drive *S. aureus* towards extinction. However, when found *in vivo*, both species can co-exist. Previous work has noted that this may due to altered gene expression or mutations. However, little is known about how the growth environment could influence co-existence of both species. Using a combination of mathematical modeling and experimentation, we show that changes to bacterial growth and metabolism caused by differences in the growth environment can determine final population composition. We found that changing the carbon source in growth medium affects the ratio of ATP to growth rate for both species, a metric we call absolute growth. We found that as a growth environment increases absolute growth for one species, that species will dominate the co-culture. This is due to interactions between growth, metabolism and metabolism altering virulence factors produced by *P. aeruginosa*. Finally, we show that the relationship between absolute growth and final population composition can be perturbed by altering the spatial structure in the community. Our results demonstrate that differences in growth environment can account for conflicting observations regarding the co-existence of these bacterial species in the literature, and may offer a novel mechanism to manipulate polymicrobial populations.

## Introduction

Microbes rarely exist in isolation. It is more common to find them as part of diverse polymicrobial communities comprised of multiple microbial species (e.g., (1)). Interactions within these communities can be synergistic or antagonistic. While cooperation can augment the stability of the community (2), competition amongst community members can stimulate the expression of gene products in an effort to drive competitors towards extinction (3). In both cases, these interactions can produce novel behaviors that are not otherwise observed when community members are grown in isolation. On the one hand, they can have beneficial functions; they can produce beneficial products, such as biofuels (4), and degrade harmful environmental pollutants (5). On the other hand, they can produce behaviors detrimental to human welfare including augmented virulence (6) and increased antibiotic resistance (7). Accordingly, understanding the general principles that shape the interactions and co-existence of members within a polymicrobial community has wide ranging implications, such as enhancing the production of beneficial products, and strategies to attenuate antibiotic resistance.

One polymicrobial community that is commonly observed in the clinic and is of increasing concern owing to the emergence of antibiotic resistance strains is composed of *Pseudomonas aeruginosa* and *Staphylococcus aureus*. These bacteria co-colonize burns (8), individuals with cystic fibrosis (9), and chronic infection sites (10, 11). Their co-occurrence leads to significantly worse healthcare outcomes (12, 13) including increased inflammation during pneumonia (9) and increased wound healing time (11, 14, 15). The interactions between *S. aureus* and *P. aeruginosa* are well understood and, at a fundamental level, influence the growth and metabolism of both species (Fig. 1A). Their co-culture induces interspecies competition, which enhances antibiotic resistance and virulence. Production of N-acetylglucosamine and autoinducer-2 from *S. aureus* augments the virulence of *P. aeruginosa* by upregulating expression of secreted virulence factors, including 4-hydroxy-2-heptylquinoline N-oxide (HQNO)(16-20). HQNO, along with the siderophores pyoverdine and pyochelin, interferes with oxidative respiration in *S. aureus* (21-24), which reduces its metabolism. This ultimately causes *S. aureus* to enter into facultative respiration leading to increased antibiotic tolerance (25). Without a fast growing competitor, and with additional nutrients released by *S. aureus* through lysis (26), *P. aeruginosa* can increase its growth and density (Fig. 1A). Overall, co-occurrence of *P. aeruginosa* and *S. aureus* alters growth and metabolism, which ultimately results in a highly virulent and antibiotic resistant infection.

**Figure 1:**
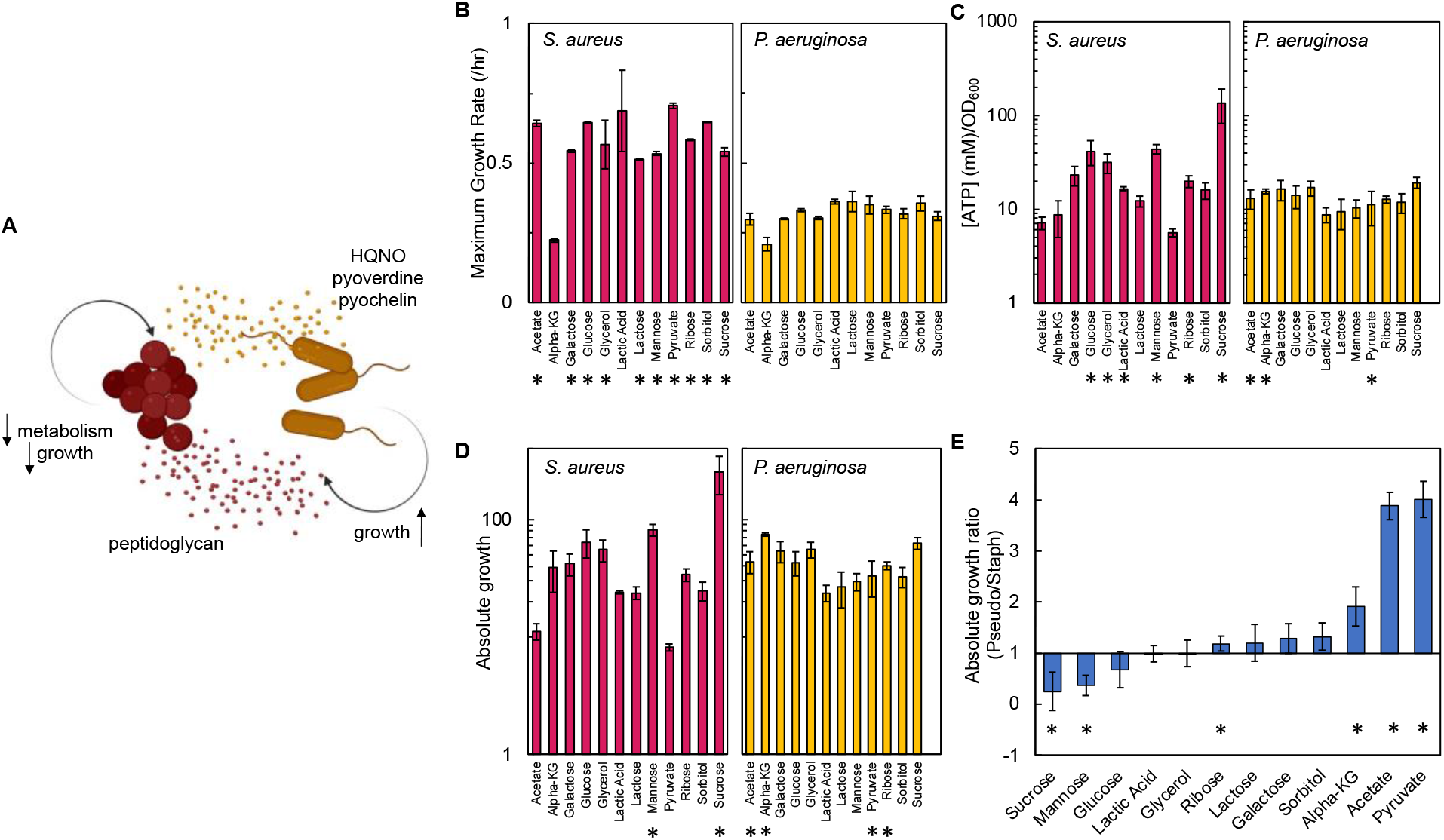
Carbon source in the growth medium affects growth and metabolism in *P. aeruginosa* and *S. aureus*; together this determines absolute growth. A) Core interactions that affect growth and metabolism of *S. aureus* (red) and *P. aeruginosa* (yellow) when in co-culture. Peptidoglycan from *S. aureus* activates the expression of virulence factors from *P. aeruginosa*. These virulence factors, pyoverdine, pyochelin and HQNO, reduce metabolism and growth of *S. aureus*, which provides a benefit to *P. aeruginosa*. B) Maximum growth rate of *S. aureus* (left) and *P. aeruginosa* (right) when grown in medium with different carbon sources. Raw growth curves in Supplementary Fig. 1A. Average from a minimum of three biological replicates. Error bars are standard deviation. * indicates a significantly greater maximum growth rate (two-tailed t-test, P < 0.05). C) The concentrations of ATP (μM) produced by *S. aureus* (left) and *P. aeruginosa* (right) grown in medium with different carbon sources. Concentration of ATP determined using a standard curve (Supplementary Fig. 1B). Standard deviation from a minimum of five biological replicates each consisting of three technical replicates. * indicates a significantly greater concentration of ATP (two-tailed t-test, P < 0.05). D) Absolute growth of *S. aureus* (left) and *P. aeruginosa* (right) grown in medium with different carbon sources. Absolute growth calculated using data shown in panels B and C. * indicates a significant difference in absolute growth between both species (two-tailed t-test, P < 0.05). E) Ratio of absolute growth between *P. aeruginosa* and *S. aureus*. Data from panel D. * indicates a significant difference as determine using data in panel D.

Previous studies have reported the co-existence dynamics of *S. aureus* and *P. aeruginosa* in human hosts. *S. aureus* is the predominant bacterial species isolated from pediatric patients with cystic fibrosis. However, during adolescence and into adulthood, the frequency of *S. aureus* decreases, while the frequency of *P. aeruginosa* increases (24, 27). One leading hypothesis to explain this trend is that *P. aeruginosa* outcompetes *S. aureus* through the production of HQNO and siderophores. Indeed, when co-cultured *in vitro, P. aeruginosa* has been shown to competitively exclude *S. aureus* over 24 hours (24). However, long term co-existence of these species has been observed in infected hosts, which has led prompted investigations into mechanisms that can account for these conflicting observations. For example, overproduction of alginate by *P. aeruginosa* (28, 29), upregulation of super-oxide dismutase (30), and decreased production of HQNO (31) are expected to facilitate co-existence. Differences in growth environments have also been suggested to play a role in facilitating co-existence. These include differences in environmental albumin concentration (32) and the production of proteins by the immune system (33). Interestingly, previous work has demonstrated that differences in nutrient availability in the growth environment can affect growth and metabolism of bacteria (34). Indeed, the presence of different nutrients and metabolites in host microenvironment has been previously shown to impact metabolism and growth of bacteria (e.g.,(35, 36)). Thus, it is possible that differences in nutrients that define the growth environment can affect the co-existence of *P. aeruginosa* and *S. aureus*. A growth environment that provides for a high metabolic rate in *S. aureus* could potentially buffer it against HQNO/siderophore driven reductions in metabolism and growth. In turn, this could facilitate co-existence and perhaps allow *S. aureus* to outcompete *P. aeruginosa*. Alternatively, growth environments that reduce growth and metabolism of *S. aureus* could make that population more sensitive to HQNO/siderophores, thus allowing dominance of *P. aeruginosa* and reducing co-existence. While these hypotheses are plausible, they have yet to be investigated. Thus, we ask can growth environment driven changes in growth rate and metabolism affect the co-existence of *P. aeruginosa* and *S. aureus* in co-culture. Addressing the role of growth environment on species interactions has implications in the bottom-up design of polymicrobial communities and may lead to novel treatment strategies in the clinic.

## Results

### Carbon source defined growth environments affects the final ratio of P. aeruginosa to S. aureus

First, we used a microplate reader to produce growth curves that allowed us to separately quantify growth rate of *P. aeruginosa* and *S. aureus* in the presence of 12 different carbon sources. We chose to use different carbon sources to perturb growth and metabolism in *P. aeruginosa* and *S. aureus* as each species has different carbon source preferences. For example, glucose is a preferred carbon source for *S. aureus*, which results in fast growth and high metabolism (37). Alternatively, *P. aeruginosa* prefers organic acids or amino acids and grows slower in glucose coinciding with a reduction in metabolism (38). We observed that in almost all cases, with the exception of α-ketoglutarate and lactic acid, *S. aureus* had a faster maximum growth rate as compared to *P. aeruginosa* (Fig. 1B). Next, we quantified the concentration of ATP produced during mid-log phase and in the presence of different carbon sources using a bioluminescent assay. We note that ATP has been previously shown to be a strong correlate of other measures of metabolism, including oxygen consumption rate and the NAD^+^/NADH ratio (39). We observed a range of concentrations of ATP (Fig. 1C). For example, when grown in medium containing glucose, glycerol, lactic acid, mannose, ribose and sucrose, *S. aureus* had a significantly higher concentration of ATP compared to *P. aeruginosa. P. aeruginosa* had a significantly higher concentration of ATP when grown in the presence of acetate, α-ketoglutarate, and pyruvate. For carbon sources of galactose, lactose, and sorbitol there was not a significant difference in the amount of ATP between the species.

To examine the combined influence of both growth and metabolism on the co-existence of both species we developed a metric called absolute growth, which we define as the ratio of ATP (mM) to growth rate (/hr). Examining the combined influence of both growth and metabolism is important as together they may impact co-existence of both bacteria. For example, consider a carbon source that provides high metabolism and slow growth to *S. aureus*. Here, *S. aureus* may be buffered against the effects of HQNO, pyoverdine and pyochelin, but may be outcompeted by *P. aeruginosa* owing to its slow growth. Alternatively, if a carbon source provides both fast metabolism and growth to *S. aureus*, it is likely to outcompete *P. aeruginosa*. Importantly, previous work has demonstrated that growth and metabolism are not linearly correlated and that nutrient availability can alter the relationship between these two variables (39-41). We found that, when provided with sucrose or mannose in the growth medium, *S. aureus* had a greater absolute growth relative to *P. aeruginosa*. Conversely, when acetate, α-ketoglutarate, ribose, or pyruvate were provided in the growth medium, *P. aeruginosa* had a greater absolute growth. Providing galactose, glucose, glycerol, lactic acid, lactose or sorbitol did not result in a statistically significant difference in absolute growth between both species.

To directly compare absolute growth for a given carbon source and between both species, we determined the ratio of absolute growth of *P. aeruginosa* to *S. aureus*. Ratios > 1 indicate that *P. aeruginosa* has a greater absolute growth, whereas ratios < 1 indicate that *S. aureus* has a greater absolute growth. We observed a range of absolute growth ratios (Fig. 1E). *S. aureus* had significantly higher absolute growth when sucrose or mannose was supplied in the growth medium. Alternatively, *P. aeruginosa* had significantly higher absolute growth when ribose, acetate, α-ketoglutarate, or pyruvate was included in the growth medium. Finally, we observed no significant difference in this ratio when galactose, glucose, glycerol, lactic acid, lactose or sorbitol was included in the growth medium. Overall, changes in carbon source provided in the growth medium alter both growth rate and metabolism leading to differences in absolute growth between both bacterial species.

### Differences in absolute growth affect the final population composition of S. aureus and P. aeruginosa

To investigate how absolute growth impacts the co-existence of *S. aureus* and *P. aeruginosa*, we chose six representative carbon sources that spanned the range of observed absolute growth values. Thus, we chose carbon sources that favored *S. aureus* (sucrose), *P. aeruginosa* (pyruvate, ribose, α-ketoglutarate) or neither (lactic acid, glucose). We initiated a co-culture of both bacterial species with equal initial densities and grew them for 24 hours without shaking at 37°C. Final bacterial density in colony forming units (CFU)/mL was measured using selective plating; cetrimide plates were used to select for *P. aeruginosa* and mannitol salt agar plates were used to select for *S. aureus*. We then determined the final density ratio of *P. aeruginosa* to *S. aureus*; values greater than one would indicate dominance of *P. aeruginosa* in the growth environment. Conversely, values less than one would indicate dominance of *S. aureus* in the growth environment. We observed that differences in the absolute growth ratio determined the final population composition; as the ratios of absolute growth increased, thus increasingly favoring *P. aeruginosa*, the final ratio of *P. aeruginosa* to *S. aureus* significantly increased (Fig. 2A). This finding was consistent when we used a weighted least squares analysis (WLS), which considers error along the y-axis when performing a regression (P = 0.0002). We also found significant differences amongst the final density ratio (P = 0.0134 (Kruskal-Wallis, Shapiro-Wilk for normality, P < 0.0001)). We did not find a significant relationship between the final ratio of *P. aeruginosa* to *S. aureus* and ATP or growth rate of either species when plotted independently (Supplementary Fig. S2). Similarly, we did not find a significant relationship between the final ratio of *P. aeruginosa* to *S. aureus* and the ratio of growth rates (growth rate *P. aeruginosa*/growth rate *S. aureus*, Supplementary Fig. S2). However, we did find a significant, albeit weaker (R= 0.7), relationship between the final ratio of *P. aeruginosa* to *S. aureus* and the ratio of [ATP] ([ATP] *P. aeruginosa*/[ATP] *S. aureus*, Supplementary Fig. S2).

**Figure 2:**
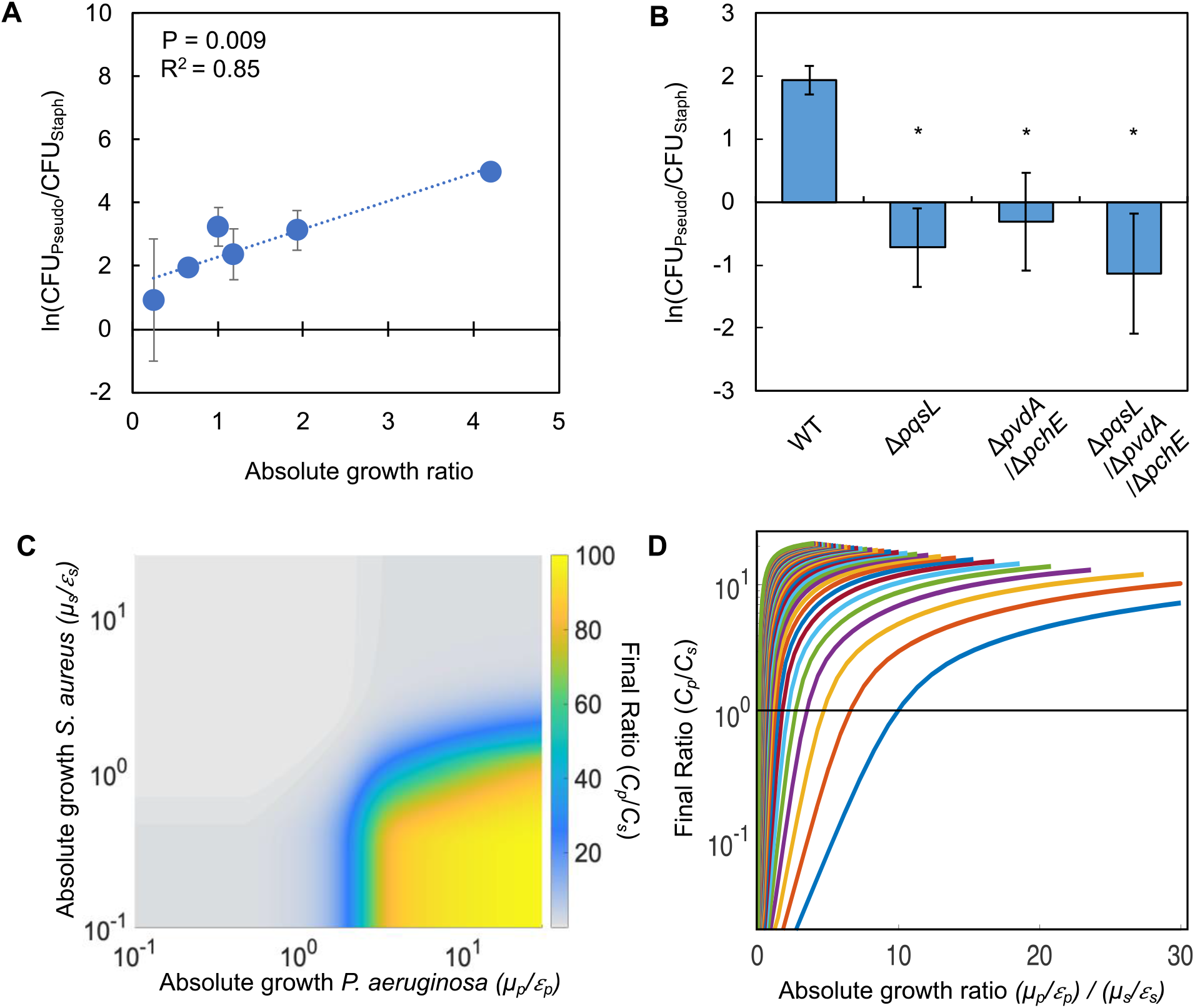
Differences in absolute growth determine the final densities of *S. aureus* and *P. aeruginosa* in co-culture. A) Final density ratio of *P. aeruginosa* to *S. aureus* after 24 hours of growth in co-culture with different carbon sources affording different absolute growth ratios. Standard deviation from a minimum of three biological replicates. R^2^ and P value shown on plot are from a linear regression. Weighted least squares regression: R^2^ = 0.98, P = 0.0002. Kruskal Wallis, P = 0.0134 (Shapiro Wilk < 0.0001). For panels A and B, final cell density in CFU for both strains show in Supplementary Fig. 2. Linear regressions between the final density ratio and [ATP], maximum growth rates, the ratio of ATP and the ratio growth rates shown in Supplementary Fig. S2. B) Final density ratio of *P. aeruginosa* knockout strains to *S. aureus* after 24 hours of growth in co-culture. Carbon source included in the growth medium was glucose. Standard deviation from a minimum of three biological replicates. * P = 0.0369 as compared to wildtype *P. aeruginosa* strain using a Mann Whitney test (Shapiro Wilk for normality < 0.001). C) Heat map showing the effect of absolute growth (*μ/ε*) of *P. aeruginosa* (*C*_*p*_) and *S. aureus* (*C*_*s*_) on the final ratio of the strains. For these simulations, growth rate (*μ*) was held constant while metabolism (*ε*) was varied. For simulations where growth rate is varied and metabolism is held constant, see Supplementary Fig. 3. For panels C and D, simulations performed using Eq. 1-3. Total simulation time = 24 hours. Parameters in Tables S2. Model description and development in *Methods*. Sensitivity analysis in Supplementary Fig. 3. D) Representative simulations showing the relationship between the ratio of absolute growth and the final density ratio of *P. aeruginosa* to *S. aureus*. For these simulations, growth rate (*μ)* and metabolism (*ε*) for *P. aeruginosa* were varied while they were fixed for *S. aureus*. Each colored line presents a combination of *μ* and *ε* for *P. aeruginosa*. Simulations using fixed values for *P. aeruginosa* and varied values for *S. aureus* shown in Supplementary Fig. 3.

To confirm that the strong relationship between the final ratio of *P. aeruginosa* to *S. aureus* and absolute growth was attributed to the production of virulence factors by *P. aeruginosa* that perturb metabolism and growth, we acquired knockout strains which lack the ability to synthesize HQNO (Δ*pqsL*), pyoverdine and pyocyanin (Δ*pvdA*/Δ*pchE*), and all three virulence factors (Δ*pqsL*/Δ*pvdA*/Δ*pchE*). When these knockout strains were co-culture with *S. aureus* in the presence of glucose as the carbon source, there was a significant increase (P = 0.0369 (Mann Whitney compared to wildtype, Shapiro Wilk; P < 0.0001). P values are henceforth only reported in the figure legends and/or supplementary data) in the final density of *S. aureus* as compared to co-culture with wildtype *P. aeruginosa* (Fig. 2B). This resulted in a decrease in the ratio of *P. aeruginosa* to *S. aureus* as compared to wild type. Overall, this indicated that the production of HQNO, pyoverdine and pyochelin plays a pivotal role in determining the interactions between *S. aureus* and *P. aeruginosa* in our experimental setup.

Towards understanding why differences in absolute growth affected the co-existence of *P. aeruginosa* to *S. aureus*, we created a simple mathematical model consisting of three ordinary differential equations. The model considers the production of virulence factors (*vir*) by *P. aeruginosa* that reduce growth and metabolism, and the growth and death of both species independently (see *Methods* for equations, model development and parameter estimation; see Supplementary Table S2 for parameters). Production of virulence factors is modeled as a modified Hill Equation that is scaled based on the density of *P. aeruginosa*. The growth of both bacterial species follows a modified logistic growth equation where growth is a product of basal growth rate, *μ*, and metabolism, *ε*. In general, *ε* approximates a maintenance coefficient, which refers to the amount of ATP that is not directly involved in generating biomass. The equation governing the growth of *S. aureus* is modified to account for the effect of virulence factors produced by *P. aeruginosa*. Here, *ε* is scaled by the amount of virulence factors (*vir*) is the system; higher concentrations of virulence factors reduce the value of *ε*. This serves to reduce the entire growth term, thus reducing the rate at which *S. aureus* grows. Consistent with our experimental definition, absolute growth for each species can be determined by calculating *ε*/*μ*. Consistent with our experimental results, our model predicts that, over a wide range of value of *μ* and *ε*, increasing the ratio of absolute growth leads to an increase in the density of *P. aeruginosa* relative to *S. aureus* (Fig. 2C-D, Supplementary Fig. S3).

### The ability of S. aureus to metabolically buffer against virulence factors is dependent upon absolute growth and the initial population composition

To gain intuition into these findings, we sought to understand how altering the initial ratio of *P. aeruginosa* and *S. aureus* would affect their final ratio after 24 hours of co-culture. Previous work has indicated that the initial population composition can impact the long term co-existence and composition of microbial communities (42). Moreover, the ability to predict the formation of microbial communities based on early conditions can allow the prediction of long term composition of microbial populations (43, 44), which has implications in the rationale design of such communities and in infectious disease.

First, we used our mathematical model to simulate the effect of altering the initial ratio of both species. Our model predicts that, for a given absolute growth ratio, as the initial fraction of *P. aeruginosa* increases, the final population density of *P. aeruginosa* also increases (Fig. 3A). Our model also predicts that, for a given initial density of *P. aeruginosa*, increasing the absolute growth ratio increases the final density of *P. aeruginosa*. Interestingly, for a given absolute growth ratio, our model predicts an initial fraction of *P. aeruginosa* that serves as a tipping point; if the initial fraction of *P. aeruginosa* exceeds this tipping point, *P. aeruginosa* will dominate the final population. Otherwise, if the initial fraction of *P. aeruginosa* is below this tipping point, *S. aureus* will dominate the final population. The initial fraction of *P. aeruginosa* that leads to this tipping point is determined by the absolute growth ratio; as this ratio increases, the initial fraction of *P. aeruginosa* at the tipping point decreases. In other words, as the absolute growth ratio increases, *P. aeruginosa* will dominate the final population when initiated at lower initial factions. We note that our model predicts that the amount of *vir* synthesized by *P. aeruginosa* increases based on both increasing initial density and an increasing absolute growth ratio (Fig. 3B).

**Figure 3:**
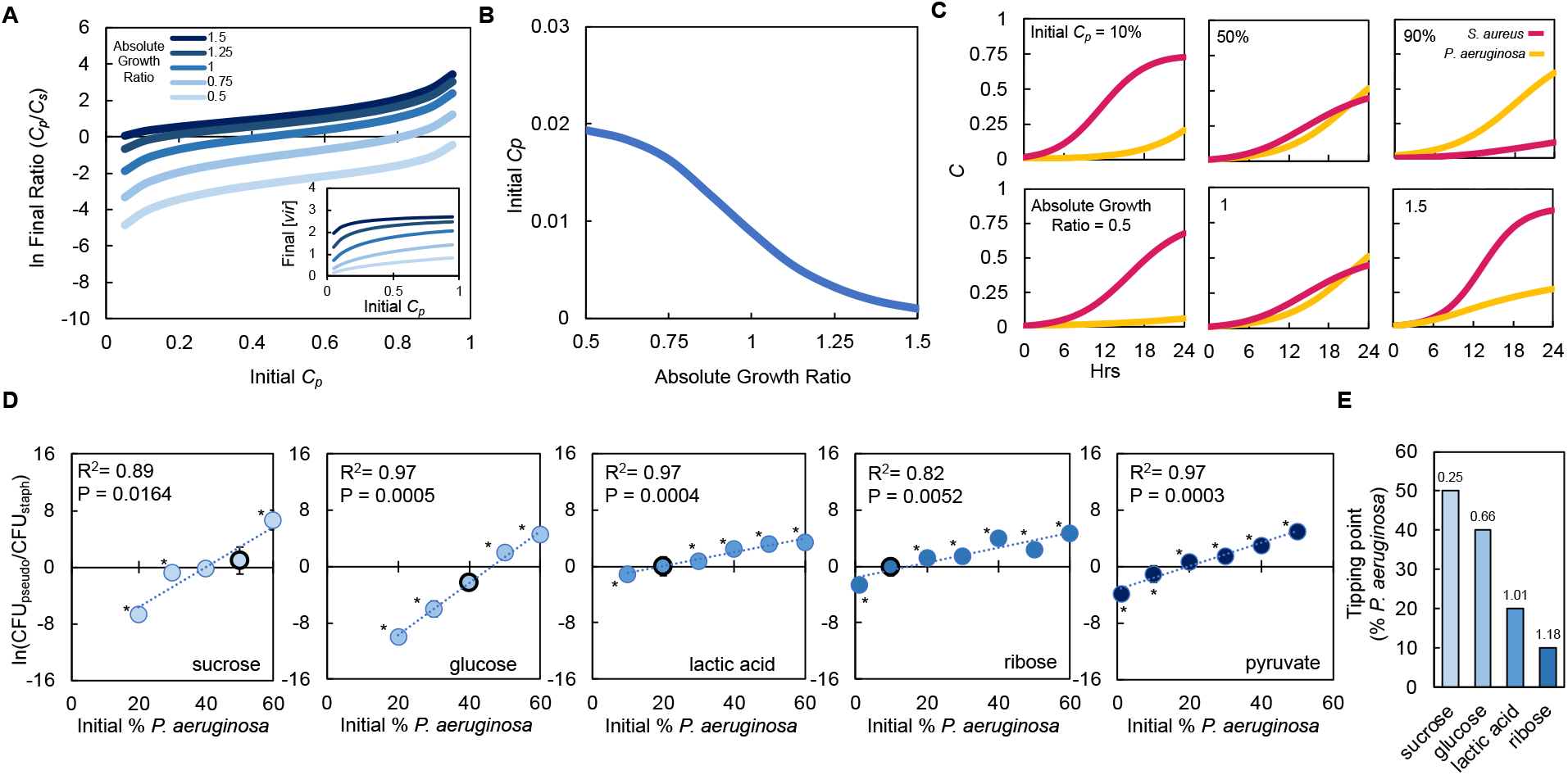
Absolute growth determines tipping point of co-existence of *P. aeruginosa* and *S. aureus* in co-culture. A) Simulations showing the effect of changing the initial fraction of *P. aeruginosa* (*C*_*p*_) and *S. aureus* (*C*_*s*_) on the final density ratio. Each colored line represents simulations performed with different absolute growth values (as indicated in the figure legend). For panels A-C, simulations performed using Eq. 1-3. Total simulation time = 24 hours. Parameters in Table S2. Model description and development in *Methods*. Inset: Simulations showing the final concentration of [*vir*] produced by *P. aeruginosa* as a function of initial density and at different absolute growth ratios. B) Tipping point. Simulations showing the lowest initial fraction of *P. aeruginosa* (*C*_*p*_) that results in a final density (*C*_*p*_/*C*_*s*_) of 1 after 24 hours as a function of the absolute growth ratio. In these simulations, metabolism (*ε*) for *P. aeruginosa* was varied but metabolism for *S. aureus* and maximum growth rates (*μ*) of both species were held constants. For simulations showing the effect of varying *ε* see Supplementary Fig. 4.. C) Simulations showing the temporal changes in the density of *P. aeruginosa* (*C*_*p*_) and *S. aureus* (*C*_*s*_). Top panels show the effect of increasing the relative initial density of *C*_*p*_ at the same absolute growth value. Bottom panels show the effect of increasing the absolute growth value while maintaining the same initial population densities (50% *C*_*p*_: 50% *C*_*s*_) D) Experimental data: The final density of a co-cultured population grown in TSB medium with different carbon sources plotted as a function of the initial percentage of *P. aeruginosa*. Standard deviation from a minimum of three biological replicates. R^2^ and P value shown on plot are from a linear regression. * indicates a significant difference in the final density of *P. aeruginosa* and *S. aureus* (Mann Whitney, P < 0.05 (Shapiro Wilk for normality < 0.0001). When all final density ratios are compared within each carbon source P < 0.007, Wilcoxon/Kruskal-Wallis (see Supplementary Table S3 for all P values). Final cell density in CFU for both strains show in Supplementary Fig. 4. Data points outlined in black indicate the tipping point; the lowest initial percentage of *P. aeruginosa* examined that led to a significantly greater amount of *P. aeruginosa* after 24 hours of growth. E) Tipping point of the population defined as the first initial percentage of *P. aeruginosa* measured where there was not a significant difference in the final density of *P. aeruginosa* and *S. aureus*. Data from panel D. Number above each bar indicates absolute growth ratio calculated using data in Fig. 1E. We did not find the tipping point for pyruvate and thus it is not included on this plot.

Our model predicts that the final population composition is largely owing to interactions between initial density, the synthesis of virulence factors that affect growth and metabolism in *S. aureus* and the ratio of absolute growth. For a given absolute growth ratio (e.g., 1), if the initial fraction of *S. aureus* (*C*_*s*_ = 0.9) is much greater than *P. aeruginosa* (*C*_*p*_ = 0.1), the amount of virulence factors produced by *P. aeruginosa* is insufficient to reduce the growth of *S. aureus* (Fig. 3C, top left panel). Coupled with a higher initial starting density, this allows *S. aureus* to outcompete *P. aeruginosa*; *S. aureus* dominates the final population. As the initial fraction of *P. aeruginosa* is increased, additional virulence factors are produced. Once *P. aeruginosa* (*C*_*p*_ = 0.5) reaches an initial fraction that is sufficiently high to produce enough virulence factors to reduce the growth of *S. aureus* (*C*_*s*_ = 0.5), *P. aeruginosa* dominates the population after 24 hours of growth (Fig. 3C, top center panel). Finally, if the initial fraction of *P. aeruginosa* (*C*_*p*_ = 0.9) is much greater than *S. aureus* (*C*_*s*_ = 0.1) the growth of *S. aureus* is sufficiently hampered such that *S. aureus* does not grow appreciably and *P. aeruginosa* dominates the population (Fig. 3C, top left panel).

Our model predicts that, for a given initial density ratio (*C*_*p*_=*C*_*s*_=0.5), increasing the absolute growth ratio benefits the growth of *P. aeruginosa*. If the absolute growth ratio is sufficiently small (0.5), *P. aeruginosa* is unable to synthesize a high amount of virulence factors and grows slower than *S. aureus*. Slow growth of *P. aeruginosa* coupled with a low amount of virulence factors synthesized allows *S. aureus* to dominate the population (Fig. 3C, bottom left panel). As the absolute growth ratio increases, it serves to benefit *P. aeruginosa*. At an intermediate absolute growth ratio (1), the amount of virulence factors produced is sufficient to reduce the growth rate of *S. aureus* while enhancing the growth rate of *P. aeruginosa*. Combined, this allows *P. aeruginosa* to dominate the population (Fig. 3C, bottom center panel). Finally, when the absolute growth ratio is large (1.5), a high amount of virulence factors is produced, which substantially reduces the growth of *S. aureus*. The increase in absolute growth ratio serves to further increase the growth rate of *P. aeruginosa*. Together, this allows *P. aeruginosa* to significantly outcompete *S. aureus* (Fig. 3C, bottom right panel).

To confirm these modeling predictions, we grew *P. aeruginosa* and *S. aureus* in co-culture as described above. We varied the initial percentage between both species and used five of the same carbon sources as used previously (Fig. 2A). We observed that our experimental results match the qualitative predictions made by our model. First, we observed that, for all absolute growth ratios tested, as the initial percentage of *P. aeruginosa* increased, the final population density was increasingly dominated by *P. aeruginosa*, which resulted in a greater final density ratio (Fig. 3D). If the initial percentage of *P. aeruginosa* was sufficiently small, *S. aureus* could dominate the population, leading to a smaller final density ratio. Moreover, as predicted by our model, we found that, as the ratio of absolute growth increased, the first initial percentage of *P. aeruginosa* measured where neither *P. aeruginosa* nor *S. aureus* dominated the population after 24 hours (or the tipping point) decreased (Fig. 3D and E).

For example, when grown in the presence of sucrose, of which *S. aureus* has a greater absolute growth, the initial percentage of *P. aeruginosa* required such that it could dominate the population after 24 hours of growth was 60% (Fig. 3E). Conversely, when grown in pyruvate, where *P. aeruginosa* has a greater absolute growth, the threshold initial percentage required for *P. aeruginosa* to dominate the population after 24 hours of growth was 10%, considerably smaller than sucrose (Fig. 3E). Overall, both our model and experimental analysis indicate that interactions between initial population composition, the synthesis of virulence factors and absolute growth determined population composition after 24 hours of growth.

### Spatial overlap and absolute growth affect co-existence of P. aeruginosa and S. aureus

Previous work has shown that relative distribution of bacteria that interact through small diffusible molecules or proteins, such as HQNO, can influence population dynamics (45). In a stationary environment, if bacteria are positioned close together in a stationary environment, the effective concentration of a small diffusible molecule sensed by the bacteria will be higher than if bacteria are positioned farther apart. Conversely, if the distribution of bacteria in the environment is homogenous, such as in a continuously shaken condition, the concentration of small diffusible molecules sensed will be roughly equal across the population. Furthermore, it has been previously shown that periodically disturbing the spatial structure of a bacterial population using a physical force not only serves to disrupt the distribution of bacteria, but can also disrupt access to small diffusible molecules (46, 47). Given that diffusible virulence factors HQNO, pyoverdine and pyochelin influence the interactions between *P. aeruginosa* and *S. aureus*, we sought to determine how the distribution of bacteria, coupled with differences in absolute growth, determined the final population composition.

To understand the relationship between the distribution of bacteria, differences in absolute growth, and their impact on population composition, we modified our series of ODEs to include a spatial overlap term, *δ*, which scales the amount of virulence factor (*vir*) sensed by *S. aureus*. High values of *δ* indicate a high degree of spatial overlap between the two populations; decreasing *δ* indicates reduced overlap, which diminishes the amount of virulence factors sensed by *S. aureus*. Experimentally, estimating *δ* would require the tracking on the positioning of multiple bacteria over both space and time whereupon a spatial overlap value could be extracted. Moreover, any movement of the plate for examination via microscopy would serve to disrupt the distribution of the cells, thus obscuring the results. Due to these challenges, we took a modeling approach to estimate trends in *δ*. We used an agent-based model (ABM, see *Methods)* to estimate the extent of overlap of the two bacterial species. The simplified ABM consists of two populations of agents that are randomly placed in a world. The agents grow and die according to logistic growth. The agents also undergo a displacement event where the spatial positioning of the agents is perturbed a given distance (called distance travelled). Increasing the number of displacement events per hour increases the degree to which the spatial positions of the agents are perturbed each hour of the simulation. In the absence of any disturbance events, the populations of bacteria do not move, which represents a stationary condition. When the number of disturbance events is very high, the population becomes well-mixed, which approximates a continuously shaken culture. An intermediate value of disturbance events would represent a populations whose spatial overlap that is perturbed periodically. After every hour of simulation using the ABM, we measure the extent of spatial overlap of the populations frequency of cohabited patches minus the expected frequency of cohabited patches (frequency of patches inhabited by *P. aeruginosa* agents x frequency of patches inhabited by *S. aureus* agents). We call this metric ‘cohabitation.’ A negative cohabitation value means populations spatially overlap less than expected based on random chance; a positive cohabitation value means that population overall more than expected. The cohabitation variable is calculated every hour, which is then averaged across the 24 hour *in silico* time period.

We observed that, over a wide range of initial population densities(Fig. 4A) and distances travelled (Supplementary Fig. 5), cohabitation followed a biphasic trend; this value first decreases with increasing disturbance events whereupon it begins to increase with increasing disturbance events. When the number of disturbance events is low (0/hr) or very high (50/hr), cohabitation is high. Towards the former, this is owing to the slow relative breakdown of their initial high degree of overlap. Towards the latter, this is due to the well mixed nature of the population where there is significant overlap between the agents. In between these two extremes the value of cohabitation is lower. This is owing to disturbance spreading-out populations and distributing cells to unoccupied areas to grow faster in the relative absence of competitors. As disturbance is infrequent, each agent spends more time in an unoccupied area over the course of the simulation, which reduces the value of cohabitation. Using the trends in cohabitation predicted by our ABM, we simulated the effect of changing *δ* and absolute growth of *P. aeruginosa* on the final density of both populations. For simplicity, we kept the absolute growth of *S. aureus* constant. We observed that at low values of absolute growth for *P. aeruginosa, δ* has little effect as *S. aureus* dominates the culture (Fig. 4B). However, at higher values of absolute growth for *P. aeruginosa*, the value of *δ* affects the final population composition. If *δ* is sufficiently large, *P. aeruginosa* dominates the population. Otherwise, if *δ* is sufficiently small, such as observed at environments that are disturbed at intermediate frequencies, *S. aureus* dominates the population.

**Figure 4:**
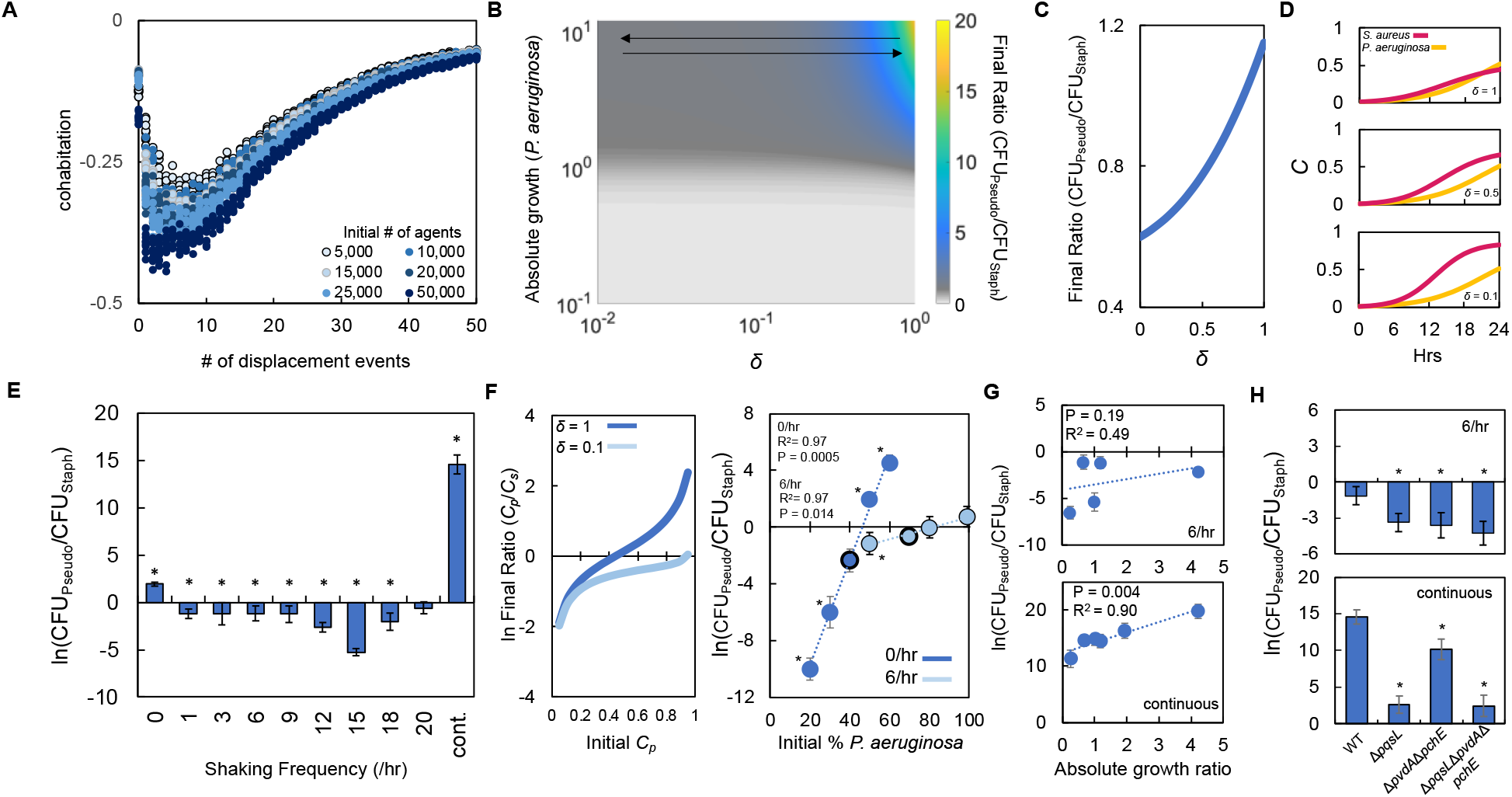
Periodically disturbing co-cultured populations of *P. aeruginosa* and *S. aureus* determines their co-existence. A) Simulation using our ABM showing the ‘cohabitation’ variable as a function of the number of disturbance events. These simulation trends were used to estimate trends in *δ* (Eq. 1-3). Each simulation was performed 10 times; each data point represents an outcome from a single simulation. See *Methods* for ABM description and parameter justification. Sensitivity analysis showing U shape trend over a wide range of parameters displayed in Supplementary Fig S5. B) Heat map showing the effect of *δ* and absolute growth of *P. aeruginosa* (fixed absolute growth of *S. aureus*) on the final ratio of *P. aeruginosa* and *S. aureus*. For panels, B-D, simulations performed using Eq. 1-3. Total simulation time = 24 hours. Parameters in Tables S2. Model description and development in *Methods*. C) Simulations showing the effect of changing *δ* on the final ratio of *P. aeruginosa* and *S. aureus* for a fixed ratio of absolute growth (1). D) Simulations showing the temporal changes in the density of *P. aeruginosa* (*C*_*p*_) and *S. aureus* (*C*_*s*_) at different values of *δ* as indicated on each panel. E) Experimental results. The ratio of *P. aeruginosa* to *S. aureus* as a function of the disturbance events in TSB medium containing glucose as the carbon source. For panels E-G, * indicates a significance difference (P < 0.05 using a Mann Whitney, Shapiro Wilk < 0.0001. P values in Supplementary Table S5) in the final density of both populations. Wilcoxon/Kruskal Wallis for all data points: P = 0.0026. Raw density data (in CFU/mL) shown in Supplementary Fig. 6. Standard deviation from a minimum of four biological replicates. F) Left: Simulations (Eq. 1-3) showing the effect of changing delta on the final population density when the initial density of *P. aeruginosa* (*C*_*p*_) is varied. Total simulation time = 24 hours. Parameters show in Supplementary Table S2. Right: Experimental data showing the final density of a co-cultured population either undisturbed (0/hr, dark blue) or disturbed at 6/hr (light blue) grown in TSB medium with different carbon sources plotted as a function of the initial percentage of *P. aeruginosa*. Standard deviation from a minimum of three biological replicates. R^2^ and P value shown on plot are from a linear regression. * indicates a significant difference in the final density of *P. aeruginosa* and *S. aureus* (Mann Whitney, P < 0.05 (Shapiro Wilk for normality < 0.0001), see Supplementary Table S6 for all P values). Wilcoxon/Kruskal Wallis for all data point within each condition P < 0.049. Final cell density in CFU for both strains shown in panel E. G) Experimental results. The ratio of *P. aeruginosa* to *S. aureus* after 24 hours of growth and with 6 disturbance events per hour (top) or in a continuously disturbed environment (bottom) in TSB medium containing different carbon sources. Difference between all data points: 6/hr: P = 0.0023 (Wilcoxon Kruskal-Waillis); continuous: P < 0.0001, ANOVA. Weighted least squares (WLS) regression analysis: 6/hr, R_2_ = 0.22 P = 0.42 ; continuous R_2_ = 0.90 P = 0.004. TSB containing glucose replotted from panel C. Raw density data (in CFU/mL) shown in Supplementary Fig. 7. Standard deviation from a minimum of three biological replicates. H) Experimental Results: The effect of 6 disturbance events per hour and continuous disturbance on the final ratio of *P. aeruginosa* knockout strains and *S. aureus* wild type strain. Raw density data (in CFU/mL) shown in Supplementary Fig. 7. Using the knockout strains resulted in a significant decrease in the ratio of *P. aeruginosa* to *S. aureus* (*, P < 0.037, Mann Whitney, P values in Supplementary Table S5). Standard deviation from a minimum of four biological replicates.

To understand these modeling predictions, we simulated temporal changes in cell density for both species independently and at three values of *δ* (Fig. 4C and D). When *δ* is small (0.1), which represents a co-culture that at an intermediate frequency (Fig. 4A), *S. aureus* grows quickly reaching a high density after 24 hours. This high growth rate is owing to a reduction in the amount of virulence factors that are encountered by *S. aureus*. At intermediate values of *δ* (0.5), which continues to represents a co-culture that at an intermediate frequency (Fig. 4A), the amount of virulence factors encountered sensed by *S. aureus* increases, which reduces its overall growth rate. However, because the growth rate of *P. aeruginosa* is sufficiently small owing to low absolute growth, *S. aureus* continues to dominate the population after 24 hours of growth. Finally, when *δ* is large (1), which represents a co-culture that is undisturbed (stationary) or very frequently disturbed (continuous shaking) the amount of virulence factors encountered by *S. aureus* is high. This significantly reduces its overall growth rate, allowing *P. aeruginosa* to outcompete it. Accordingly, *P. aeruginosa* dominates the population after 24 hours under this condition. Importantly, under each of these simulated conditions the total amount of virulence factor synthesized is not altered ([*vir*] = 1.77 μM); only the concentration sensed by *S. aureus* owing to changes in *δ*.

To test these predictions, we periodically disturbed co-cultured populations of *S. aureus* and *P. aeruginosa* using the linear shaking function of a microplate reader. We previously demonstrated that using this function could alter the spatial distribution of bacteria (46, 47). We grew both bacterial species using glucose as the carbon source as it represented an intermediate ratio of absolute growth. As above, when both bacteria species were initiated at equal percentages, the use of glucose allowed *P. aeruginosa* to dominate the population in the stationary condition. However, as predicted by our model, *S. aureus* dominated the final population composition over a range of intermediate disturbance frequencies (1/hr-18/hr, Fig. 4E). However, as the disturbance frequency increased outside of this range, there was no statistical difference between the final density of both populations and the density of *P. aeruginosa* was significantly greater than *S. aureus*. We note that *S. aureus* was observed to dominate the population at a disturbance frequency of 6/hr at lower initial dilutions of the population (Supplementary Fig. S6). Moreover, we also found that disturbance at 6/hr shifted the tipping point of the population to a higher initial percentage of *P. aeruginosa* as compared to the stationary condition (0/hr, Fig. 4F). This finding was consistent both *in silico* (Fig. 4F, left panel) and *in vitro* (Fig. 4F, right panel).

Next, we investigated how changing absolute growth through the use of different carbon sources might impact the ability of *S. aureus* to dominate at intermediate, but not high, disturbance frequencies, such as a continuously mixed population. Accordingly, we grew co-cultured populations of *S. aureus* and *P. aeruginosa* in four additional carbon sources that spanned the range of growth efficiencies and disturbed these populations at a disturbance frequency of 6/hr or continuously. As predicted by our model, at a disturbance frequency of 6/hr, *S. aureus* dominated the population after 24 hours of growth regardless the absolute growth ratio (Fig. 4G, top panel). Under continuous shaking, we observed that as the ratio of absolute growth increased, the final ratio of *P. aeruginosa* and *S. aureus* increased (Fig. 4G, bottom panel). We note that we did not find a significant relationship between the final ratio of P. aeruginosa to S. aureus (at both 6/hr and continuous disturbance) and the ratio of growth rates (*P. aeruginosa* growth rate/*S. aureus* growth rate or the ratio of [ATP] (P. *aeruginosa* [ATP]/*S. aureus* [ATP]).

In both the 6/hr and continuous disturbance conditions, metabolism altering virulence factors produced by *P. aeruginosa* continues to have a predominant role in mediating the interaction between both bacterial species. When *S. aureus* was co-cultured with knockout strains of *P. aeruginosa* (Δ*pqsL*; Δ*pvdA*/Δ*pchE*); Δ*pqsL*/Δ*pvdA*/Δ*pchE*), resulted in an increase in the final density of *S. aureus* relative to *P. aeruginosa* when disturbed at 6/hr (Fig. 4H, left panel) or continuously (Fig. 4H, right panel). Overall, our simulations and experiments suggest that access to metabolite altering virulence factors driven by spatial organization and absolute growth can determine the final population composition of *P. aeruginosa* and *S. aureus*.

## Discussion

Through a series of empirical and analytical investigations, we have shown that differences in absolute growth, a combined metric that encompasses the relationship between growth rate and metabolism, can determine the population composition of *S. aureus* and *P. aeruginosa*. In a stationary environment, as the absolute growth ratio increases, the final density ratio of *P. aeruginosa* and *S. aureus* increases. While this is also true in a continually disturbed environment, this relationship falls apart in a periodically disturbed environment. Our model and experiments suggest that this is owing to changes in access to metabolism altering virulence factors produced by *P. aeruginosa*; in stationary or continuously disturbed environments, the spatial overlap of both bacteria is predicted to be high, which facilitates interactions between these virulence factors and *S. aureus*. However, in periodically disturbed environments, the overlap between bacteria is reduced; this limits interactions between *S. aureus* and virulence factors. *S. aureus* then dominates the population. Absolute growth appears to be the most accurate metric within the scope of our study to quantify the changes in the final ratio between *P. aeruginosa* and *S. aureus*. While we did find a significant relationship between the final ratio of both species and the ratio of ATP of both species when grown in the stationary condition, the relationship was not as strong as growth efficiency (R^2^ = 0.85 (absolute growth) vs 0.70 ([ATP] ratio)). Moreover, we did not find a significant relationship between the final ratio of both species and the ratio of ATP when grown in the continuously disturbed condition.

Previous work has shown that niches in the human body, which can vary in environmental conditions, can promote bacterial growth and persistence (48). Spatiotemporal differences in nutrient availability within these niches can promote the growth of different bacteria (34, 49). Even within the same niche, such as sputum isolated from individuals with cystic fibrosis, the concentration of critical nutrients, such as amino acids and carbon sources, can differ from host to host (50). Variations in nutrient availability within or between niches may alter growth and metabolism of the resident bacteria, and may serve to perturb interactions within polymicrobial communities. In the case of *P. aeruginosa* and *S. aureus*, difference in nutrient availability in sputum (50) or the epithelium (51) may alter growth, metabolism and the interactions between these species. This could alter their co-existence dynamics leading to host and/or niche specific colonization dynamics. Differences between *in vitro* and *in vivo* growth and metabolism owing to the nutritional composition of the growth environment may also help account for differences in the co-existence dynamics reported previously (24, 27).

It has been previously recognized that spatial organization facilitated by an undisturbed growth environment can facilitate the co-existence of bacteria, including *P. aeruginosa* and *S. aureus* (52). Importantly, the production of HQNO from *P. aeruginosa* can impact the pattern of spatial organization of this co-cultured population (52). While we did not explicitly quantify spatial organization experimentally, in theory, co-culture in an undisturbed condition should facilitate the creation of spatial organization. We found that, in this environment, the absolute growth ratio could predict the final population composition; increasing the absolute growth ratio increased the final density of *P. aeruginosa* relative to *S. aureus*. When disturbed periodically using a physical force, *S. aureus* dominated the final population composition. However, a clear trend between the absolute growth ratio and the final population composition was not observed. Our model suggests that these disturbances decrease the spatial overlap of both populations, and limit the amount of virulence factors that are sensed by *S. aureus*. Importantly, physical forces that are encountered where bacteria can infect hosts, including luminal areas(53, 54), can fluctuate, which could serve to periodically alter the spatial positions, or overlap, between both population. Thus, the degree to which a population of *S. aureus* and *P. aeruginosa* is disturbed may influence their ability to interact; this would impact the population composition. Importantly, we cannot rule out that increasing the amount of disturbances might have also impacted oxygenation of the medium, or caused unknown transcriptional responses in the bacteria. However, our model, and experiments using knockout strains, suggest that co-existence of *S. aureus* and *P. aeruginosa* in periodically disturbed environments is largely due to interactions governed by HQNO, pyoverdine and pyochelin. While these disturbances have been previously shown to impact bacteria cooperation (55) and the production of virulence factors (46), to our knowledge, this is the study reporting that such disturbances can alter the population composition in a polymicrobial community. Thus, these disturbances represent a new method by which the composition, and potentially the functionality, of a polymicrobial community might be perturbed.

Co-culture of *S. aureus* and *P. aeruginosa* leads to changes in antibiotic susceptibility. For example, HQNO produced by *P. aeruginosa* interferes with ATP production in *S. aureus*. This causes a reduction in the metabolism of *S. aureus* and increases its resistance to aminoglycoside antibiotics. Important, a reduction in metabolism has been previously noted to decrease antibiotic efficacy across a wide range of bactericidal antibiotics (39). Interestingly, HQNO can also increase membrane permeability of *S. aureus*, which increases the efficacy of antimicrobials such as chloroxylenol and fluoroquinolones (56). Manipulation of absolute growth using different carbon sources may serve to rationally increase the efficacy of antibiotics. For example, carbon sources that cause *S. aureus* to have a significantly greater absolute growth relative to *P. aeruginosa* would limit the growth of *P. aeruginosa*, and thus reduce the total concentration of HQNO produced. This would allow the metabolism of *S. aureus* to remain relatively high, thus promoting the efficacy of antibiotics, such as aminoglycosides. Conversely, promoting the production of *P. aeruginosa* by using carbon sources that increase its absolute growth relative to *S. aureus* would augment the production of HQNO. This could increase the membrane permeability of *S. aureus* found in biofilms, thus augmenting sensitivity to certain antibiotics, such as fluoroquinolones. Thus, rational manipulation of absolute growth in polymicrobial interactions may serve to differentially alter susceptibility to existing antibiotics.

## Methods

### Strains and growth conditions

*P. aeruginosa* strain PA14 and *S. aureus* strain RN4220 were used in this study. *P. aeruginosa* mutants Δ*pqsL*, Δ*pvdA/*Δ*pchE* and Δ*pqsL/*Δ*pvdA/*Δ*pchE* were obtained from (56). Single colonies of *P. aeruginosa* and *S. aureus* isolated from Luria-Bertani (LB) agar medium (MP Biomedicals, Solon OH) were inoculated overnight in 3 mL of liquid LB medium and shaken (250 RPM and 37°C) in 15 mL culture tubes (Genessee Scientific, Morrisville, NC). We then washed the cells in fresh tryptic soy broth (TSB) medium (Becton, Dickinson and Company, Franklin Lakes, NJ) or modified TSB medium [Soytone (Thermo Fisher Scientific, Waltham, MA), Tryptone, Dipotassium Phosphate, and Sodium Chloride (VWR International, Radnor, PA)] with the addition of a carbon source other than glucose (0.25%). Carbon sources tested in this manuscript were as follows: D(+) mannose (Acros Organics, Geel, Belgium), glycerol (Acros Organics), D(+) ribose (Acros Organics), 2-ketoglutaric acid (Alfa Aesar, Tewksbury, MA), D(+) galactose (Alfa Aesar), D-lactose (Alfa Aesar), sodium pyruvate (Alfa Aesar), sorbitol (Fisher Scientific, Waltham, MA), D-glucose (Fisher Scientific), D(+) lactic acid lithium salt (MP Biomedicals), ands sucrose (Sigma-Aldrich, St Louis, MO).

### Growth rate

Overnight cultures were washed once in modified TSB lacking a carbon source and diluted 200-fold into 200 μL of fresh TSB with a carbon source as indicated in the text. This was the placed in the wells of a 96 well plate. The medium was overlaid with 70 μL of mineral oil to prevent evaporation. Using a Perkin Elmer Victor X4 (Waltham, MA) plate reader, cell density was determined every 10 minutes (optical density at 600nm (OD_600_)) for approximately 15 hours at 37°C. OD_600_ values from cell free medium were subtracted from all measurement prior to growth curve fitting. To help remove any artifacts of background and to ensure all data was initiated at the same starting point, we log-transformed and normalized to the initial minimum density. Together, this reduces the error when performing curve fitting. Growth rate was determined by fitting a logistic curve to the data using a custom MATLAB (MathWorks Inc., Natick, MA) code (57). Average residuals for each biological replicate are shown in Supplementary Table S7; lower residual values indicate a stronger fit between our experimental data and the curve fitting.

### Determining the Concentration of ATP

Overnight cultures were washed once with modified TSB medium lacking a carbon source. Bacteria were then diluted 10-fold with TSB supplemented with 2.5% of a carbon source in a 6-well cell culture plate (Genesee Scientific). The plate was overlaid with two AeraSeal sealing membranes (Sigma-Aldrich) and shaken at 110 RPM pre-set to 37°C for 3 hours whereupon the bacteria entered log phase growth. 100 μl of these monocultures was added to the wells of an opaqued walled 96 well plate and ATP was measured using a bioluminescent kit as recommended by the manufacturer (BacTiter-Glo Microbial Cell Viability Assay, Promega, Madison, WI). An ATP standard curve was prepared using purified ATP (Sigma Aldrich). All ATP measurements were normalized to cell density (OD_600_) and convert to concentration using a standard curve prepared using purified ATP (Supplementary Fig S1B).

### Co-culture Assays

Single colonies of *P. aeruginosa* and *S. aureus* were grown overnight and washed with TSB medium. Bacteria were then diluted to an OD_600_ of 0.1, which leads to an equal initial density between both strains (Fig. S2). The normalized cells were then diluted 100-fold in 6-well cell culture plates, overlaid with two AeraSeal sealing membranes, and placed in an undisturbed incubator for 30 minutes at 37°C. For most assays, the monocultures of *P. aeruginosa* and *S. aureus* were then combined in a 1:1 ratio to a volume of 3 mL, re-sealed with AeraSeal film. We altered the initial bacterial ratio by scaling the addition of *P. aeruginosa* monoculture to the co-culture to accommodate for a total volume of 3 mL. For the stationary (undisturbed) and continuously disturbed conditions, the plates were either placed in a stationary incubator at 37°C or shaken at 110 RPM at 37°C, respectively, for 24 hours. The plates that were periodically shaken were placed in a Victor X4 plate reader pre-set to 37°C (slow setting, frequency, 10 seconds per shaking event, orbital shaking feature, radius = 5mm) at the indicated frequency. After 24 hours of growth, 1 mL of culture was removed and washed with 1x phosphate buffered saline (PBS) (Fisher Scientific). The culture was then sonicated in a water bath for two minutes to reduce clumping of *S. aureus*. We then performed a serial dilution and selective plating on mannitol salt agar (Becton, Dickinson and Company, Franklin Lakes, NJ, selects for *S. aureus*) and cetrimide (Oxoid, Basingstoke, England, selects for *P. aeruginosa*) Following 20 hours of incubation, we determined the number of colony forming units (CFU) grown on the selective agar medium.

### Statistical Analysis

Statistical analysis as indicated in the text or figure legend. Unpaired t-tests (unequal variance) were performed using Microsoft Excel (Redmond, WA). Additional tests were performed in JMP Pro 16 (SAS Institute Inc., Cary, NC). We took the natural logarithm of the ratio of *P. aeruginosa* to *S. aureus* to linearize the data; these values were used for linear regressions. To test if the ratios themselves differed within a dataset using a Mann-Whitney or Wilcoxon/Kruskal Wallis, we used the raw ratio data that was not ln transformed. When performing a weighted least squares (WLS) analysis, we first converted standard deviation to variance, and then used inverse variance to weight each data point. Thus, data points with larger standard deviations have less weight when performing a linear regression. A Shapiro Wilk test was used to assess normality.

### Mathematical modeling - ODE

We modeled the interaction between *S. aureus* and *P. aeruginosa* using three ordinary differential of virulence factors that affect metabolism in *S. aureus* (Eq. 1), growth of *P. aeruginosa* (Eq. 2), and growth of *S. aureus* (Eq. 3).

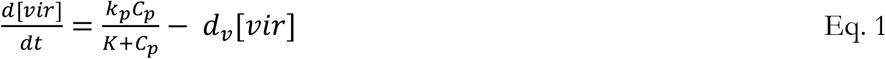

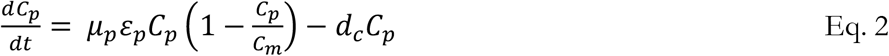

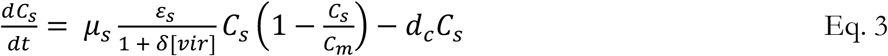

where *k*_*p*_ represents the maximum synthesis rate of virulence factors that affect metabolism in *S. aureus* (*vir*), *d*_*v*_ represents the degradation rate of virulence factors, *C*_*p*_ and *C*_*s*_ represents the density of *P. aeruginosa* and *S. aureus*, respectively, *μ*_*p*_ and *μ*_*s*_ represents the maximum growth rate of *P. aeruginosa* and *S. aureus*, respectively, *ε*_*p*_ and*ε*_*s*_ represents metabolism of *P. aeruginosa* and *S. aureus*, respectively *C*_*m*_ represents the carrying capacity of the medium normalized to 1, *d*_*c*_ represents death rate of bacteria, and *δ* represents the amount of spatial overlap in the system. The synthesis of *vir* is modeled using a modified Hill Equation and is dependent upon cell density as previous work has indicated non-linear and density dependent production of HQNO and pyoverdine (17, 58). The growth term of Eq. 2 and 3 is modeled using a simplified version of Monod’s growth. Maximum growth rate, *μ*, is scaled by the metabolic state, *ε*, of the cell; the product of these two variables determines the effective growth, which is used to inform the logistic growth term. *ε* approximates a maintenance coefficient. The growth term of *S. aureus* (Eq. 3) is scaled by the concentration of virulence factors that affect metabolism. As observed in previous work, increasing concentrations of HQNO, pyoverdine and pyochelin decrease metabolism, which in turn decreases growth rate. An increase in the concentration of these virulence factors would reduce the value of *ε*. This in turn would reduce the entire growth term (*μ* x *ε*), thus capturing the dynamics of previous experimental work. *δ* scales the concentration of virulence factors that are sensed by *S. aureus*; it is a simplification of the spatial distribution of *S. aureus* and *P. aeruginosa* during disturbance. Lower values of *δ* indicate reduced spatial overlap between *S. aureus* and *P. aeruginosa*.

### Parameter estimation -ODE

For simulations were growth rate (*μ*) was not varied, we estimated the growth rate of both bacterial species using the average growth rate found in the study (Fig. 1B, *S. aureus* = 0.57 +/- 0.12 ; *P. aeruginosa* = 0.32 +/- 0.04). We also ensured that the growth rate of *S. aureus* was greater than that of *P. aeruginosa*, which was consistent with the general trends in our experiments (Fig. 1B) The range of values of *ε* were estimated using previously published maintenance coefficients (59-61). The value of *k*_*p*_ was estimated as it represents a lumped term that captures the synthesis of three different virulence factors; pyoverdine, HQNO, and pyochelin. The average amount of pyoverdine synthesized by *P. aeruginosa* has been reported to be approximately 0.3 μM/hr (58). The approximate amount of HQNO synthesized by a population of *P. aeruginosa* is approximately 3500 μM/hr (17). Finally, the amount of pyochelin synthesize is approximately 0.002-0.02 μM/hr (62). Given the wide range of synthesis values, and the simplification of our *k*_*p*_ term, we approximated *k*_*p*_ of 1 μM/hr. We note that increasing or decreasing this value tenfold maintains the qualitative trends in our simulations (Supplementary Fig. S3). To estimate the value of *d*_*v*_, we considered previously reported degradation rates of pyoverdine (dilution rate of approximately 0.5/hr (63)) and HQNO (decay of approximately 2 mg/mL over 10 hours, or ∼0.01/hr (64)). As above, given the wide range of dilution/decay rates, we estimated a central value of 0.1/hr; increasing or decreasing this value 10-fold does not change the qualitative nature of our predictions (Supplementary Fig. S3). We estimated the order of magnitude of the cell death (*d*_*c*_) using previously published data (65). We fit the value of *K* to our experimental data. Values of *δ* were benchmarked to a value of 1, which indicates near complete spatial overlap between *P. aeruginosa* and *S. aureus*. Reducing the value of *δ* served to decrease the spatial overlap of both species, thus reducing the effect of these virulence factors. Trends in *δ* were approximated using agent based modeling (see below, Fig 4A). Simulations were performed for *t* = 24 hrs, which is the same amount of time over which our experiments occurred. The general trend of our simulations remains the same when parameters are increased or decreased 10-fold from the base set of parameter values (Supplementary Table S2).

### Mathematical modeling – agent based model

We constructed an agent-based model (ABM) in the program Netlogo (66) to better understand the amount of spatial overlap of two bacterial populations under various disturbance conditions. To coincide with our experimental approach, the model begins with equal-sized populations of *S. aureus* and *P. aeruginosa* cells (agents) placed in completely overlapping distributions at the center of the virtual world and spread-out based on a normal distribution. The virtual world is a 2-dimensional grid comprised of 10,201 patches (101 patches vertically x 101 patches horizontally). Each round, individual agents reproduce to gradually grow the population logistically (*r* = 0.5) from its initial size to reach the final carry capacity of 200,000 agents. While there are no antagonistic interactions between the populations, growth is still limited by local density as each patch has a carrying capacity (of overall carrying capacity divided by number of patches). Therefore, reproduction is diminished in crowded patches and more favorable in less occupied patches. To mimic periodic spatial disturbances, each time step all cells are displaced (called ‘distance travelled’) in random directions at a percentage distance of the virtual world (base case is 1% or one patch). These disturbances were varied from occurring once each round, to up to 50-times each round, or not at all. Simulations were iterated under given disturbance conditions for 24-rounds, to resemble 24-hour laboratory experiments. The purpose of these simulations was to observe the extent that disturbance regime alone influences the extent of overlap of competing populations. The amount of overlap was represented by the variable, cohabitation, which is the observed frequency of patches hosting agents from both population minus expected random frequency of co-habitation. Specifically, this was calculated as the number of observed co-occupied/total occupied patches, subtracted by the product of frequency of patches occupied by *S. aureus* agents and frequency of patches occupied by *P. aeruginosa*. Therefore, a positive value indicates that the two populations cohabit patches are a frequency greater than expected randomly, and a negative value indicates that individuals are less likely to co-occupy patches than expected randomly. For each simulation, this value was averages across all 24-rounds.

## Acknowledgements

This research was sponsored by the Army Research Office and was accomplished under Grant Number W911NF-18-1-0443. The views and conclusions contained in this document are those of the authors and should not be interpreted as representing the official policies, either expressed or implied, of the Army Research Office or the U.S. Government. The U.S. Government is authorized to reproduce and distribute reprints for Government purposes notwithstanding any copyright notation herein. *S. aureus* strain RN4220 was obtained through BEI Resources, NIAID, NIH. The *P. aeruginosa* mutants Δ*pqsL*, Δ*pvdA/*Δ*pchE* and Δ*pqsL/*Δ*pvdA/*Δ*pchE* were a kind gift from Dr. George A. O’Toole.

## Data sharing plan

The authors will freely share their data and modeling code by posting it to the Dryad Digital Repository upon publication of the manuscript.

